# Pupil dilation scales with movement distance of real but not of imagined reaching movements

**DOI:** 10.1101/2023.01.11.521813

**Authors:** Dimitris Voudouris, Immo Schuetz, Tabea Schinke, Katja Fiehler

## Abstract

Pupillary responses have now been reliably identified for cognitive and motor tasks, but less is known about their relation to mentally simulating movements (known as motor imagery). Previous work found pupil dilations during the performance of simple finger movements, where peak pupillary dilation scaled with the complexity of the finger movement and force required. Recently, pupillary dilations were reported during imagery of grasping and piano playing. Here we examined whether pupillary responses are sensitive to the dynamics of the underlying motor task for both executed and imagined reach movements. Participants reached or imagined reaching to one of three targets placed at different distances from a start position. Both executed and imagined movement times scaled with target distance, and they were highly correlated, confirming previous work and suggesting that participants did imagine the respective movement. Increased pupillary dilation was evident during motor execution compared to rest, with stronger dilations for larger movements. Pupil dilations also occurred during motor imagery, however they were generally weaker than during motor execution and they were not influenced by imagined movement distance. Instead, dilations during motor imagery resembled pupil responses obtained during a non-motor imagery task (imagining a previously viewed painting). Our results demonstrate that pupillary responses can reliably capture the dynamics of an executed goal-directed reaching movement, but suggest that pupillary responses during imagined reaching movements reflect general cognitive rather than motor-specific components of the motor imagery process.

**New and noteworthy:** Pupil size is influenced by the performance of cognitive tasks. Here we demonstrate that pupil size increases during execution and mental simulation of goal-directed reaching movements compared to rest. Pupil dilations scale with movement amplitude only during motor execution, whereas they are similar during motor imagery and a non-motor imagery, cognitive task.

## Introduction

When mounting a new piece of furniture or about to drive a screw into the wall, mental simulation can help to find the correct direction to rotate the hand holding the screwdriver. Imagining a movement and its sensory consequences is a common process that occurs in several daily instances, such as when learning a new motor skill or when interacting with a novel environment. This is typically called motor imagery and refers to the state during which one mentally simulates the sensations caused by one’s own movement, without actually executing this movement. Motor imagery seems to also foster actual motor related processes. For instance, motor imagery training can lead to an increase in associated muscle force (Ranganathan et al., 2004), and can lead to improved kinematic performance later on (Ruffino et al., 2017).

Motor imagery is thought to involve the same processes as those involved during the planning and execution of the real action, with the main difference being that, in motor imagery, actual motor execution is interrupted, possibly at some cortico-spinal level (Jeannerod, 1994). Claims of functional equivalence between motor imagery and execution are traditionally based on findings of the so-called mental chronometry paradigms, which assume that if executed and imaginary movements are governed by similar processes, then their temporal organization should also be similar. In support of these assumptions, the time it takes to write a sentence or to draw a shape is similar to the time it takes to mentally simulate the respective action (Decety & Michel, 1989). In addition, executed and imagined movement times are modulated by the characteristics of the underlying task. For example, they both get longer when a hand action is performed or imagined either with an additional mass on the hand (Papaxanthis et al., 2002; Munzert et al., 2015) or using the non-dominant hand (Decety & Michel, 1989). Imaginary movement times also obey predictions of Fitts’ law, as they get longer when imagining to walk through narrower than wider gates (Decety & Jeannerod, 1996), suggesting that imagined movements consider the underlying dynamics about the required precision. Yet, imaginary movement times are sometimes reported to be shorter (Calmels & Fournier, 2001) and sometimes longer (Reed, 2002) than actual movement times.

There is evidence that executed and imagined movements are equivalently represented also in physiological measures. Overlapping brain activity has been found between executed and imagined actions (Bonnet et al., 1997; Decety, 1996; Decety et al., 1990; Hashimoto & Rothwell, 1999; Lorey et al., 2011; Munzert et al., 2009), as well as between imagined and observed actions (Munzert et al., 2008) such in parietal and motor cortices as well as in the cerebellum. Motor imagery was also shown to affect pupillary responses (O’Shea & Moran, 2016; Richer & Beatty, 1985; Rozado et al., 2015; Rozado et al., 2017). Constrictions of the pupil are controlled through the sphincter muscle, innervated by the parasympathetic system, whereas pupil dilations are caused by the dilator muscle, innervated by the sympathetic system (Mathôt, 2018). Pupil size is generally sensitive to a number of psychophysiological factors, including illumination, arousal, and mental effort (for a detailed description please refer to Einhaeuser, 2017 and Mathôt, 2018), as well as to motor related factors (Bernick & Oberlander, 1968; Hupé et al., 2009). For instance, the pupil starts dilating already 1.5 s before an upcoming self-paced finger flexion, with the peak of this dilation being ∼0.5 s after movement onset (Richer & Beatty, 1985). This response scales with the number of fingers involved in the movement as well as with the required force that the finger should produce, suggesting that movement complexity and effort modulate the strength of the dilation. Similarly, pupil dilations to a cued finger movement are larger for movements that are more difficult to encode and prepare (Moresi et al., 2008a, 2008b). Pupillary responses also have higher peaks and these peaks occur later when performing joystick movements with greater difficulty in a Fitts’ Law paradigm (Jiang et al., 2014). However, pupil diameter appears smaller during joystick movement preparation and execution for targets with greater requirements on reaching precision (Fletcher et al., 2017). Pupillary responses are also evident during goal-directed movements under online motor control, such as grasping or piano playing, and recent work reported similar findings also for imagined movements (O’Shea & Moran, 2016; Rozado et al., 2015; Rozado et al., 2017). These pupillary responses seem comparable between executed and imagined movements, however this is based on averaged pupillary dilations across the time course of the pupillary response, rendering the understanding of the temporal evolution of pupillary responses rather ambiguous. In addition, it remains unclear whether pupillary responses during goal-directed movements are influenced by the specific movement dynamics, such as different levels of complexity, as shown for simple hand movements (e.g., Richer & Beatty, 1985). Accordingly, relatively little is currently known about pupillary responses to imagined movements and their temporal evolution.

In this study, we examined and compared the characteristics and temporal evolution of pupillary responses during motor execution and motor imagery of goal-directed reaching movements. First, we investigated whether specific characteristics of the pupillary response, such as peak dilation and its latency, are influenced by executed and imagined reaching movements. Second, we examined whether the temporal evolution of the associated pupillary responses is different between motor execution and motor imagery. Third, we aimed at understanding whether movements of different amplitude differentially influence the pupillary response characteristics and their temporal evolution. Participants were asked to perform a reach or an imagined reach toward targets at different distances while we recorded their executed and imaginary movement times and associated pupillary responses. We expected that both executed and imaginary movement times would be similarly modulated by target distance (Papaxanthis et al., 2002; Munzert et al., 2015). Specifically, they should both be longer for more distant targets, while executed and imaginary movement times should correlate with each other. To examine the role of motor execution and imagery on pupillary responses, we examined the modulation of (a) the peak pupil dilation following actual and imaginary movement onset, (b) its latency, as well as (c) the temporal evolution of the pupillary responses between tasks. Finally, we explored whether pupillary responses are affected by target distance. To assess possible non-motor cognitive effects of motor imagery on pupillary responses, we further examined whether the characteristics and temporal evolution of pupillary responses differ between the motor and a non-motor imagery task (i.e. imagining a previously seen painting).

Identifying pupillary correlates of motor imagery (beyond the well-established effects of cognitive load during an imagery paradigm; see e.g., Hess & Polt, 1964) would allow to determine whether participants actually imagined a movement during a given task, without requiring an additional overt response such as a button press at the end of the imagined movement. Additionally, comparable changes in pupillary dilation with movement distance for both executed and imagined reaching would provide further evidence to the recruitment of similar processes in both tasks.

## Experiment 1

### Methods

#### Participants

Thirty participants (16 women, 14 men; age range 20-33 years) volunteered for this experiment. They were all right-handed according to the German translation of the Edinburgh Handedness Inventory (range: 50-100; Oldfield, 1971), except for one participant who was ambidextrous with a handedness index of 10. Participants were free of any known muscular and neurological issues at the moment of the experiment, and received either 8 €/hour or course-credits for their participation. The experiment was approved by the local ethics committee of the Justus-Liebig University Giessen and was in accordance with the declaration of Helsinki (2013, except of §35, pre-registration).

#### Experimental Setup

Participants were seated at a table with their head resting on a chin rest. An LCD monitor (VIEWPixx, VPixx Technologies Inc, Saint-Bruno, QC, Canada) was used to present visual stimuli at a distance of 80 cm to the participant’s eyes (screen resolution 1920 × 1080 pixels, refresh rate 60 Hz). The participant’s right hand rested on the mouse button of a USB touchpad (Perixx PERIPAD-504, Conrad Electronic, Hirschau, Germany), which was used as the start button and was placed 30 cm to the right and 35 cm forward of their body midline. An infrared LED marker was affixed to the nail of their right index finger and recorded the 3D index finger position throughout each trial at a rate of 100 Hz using an Optotrak Certus device (Northern Digital, Waterloo, ON, Canada). An Eyelink II eye tracking device (SR-Research Ltd., Ottawa ON, Canada) was affixed to the participant’s head by means of a headband and recorded movements and pupil size of the right eye at a sampling rate of 500 Hz. Three target positions (3 × 3 cm) were marked on the table surface using thin masking tape. The 3 targets were at 15, 30 and 45 cm from the touchpad, along the participants’ lateral axis.

All visual stimuli were presented on a medium gray screen (18.5 cd/m²) which was kept constant throughout the experiment to avoid any influences of screen luminance on pupil size (Ellis, 1981). For the same reason, window blinds in the experiment room were kept closed and lighting was kept constant for all participants (ca. 370 lux). Instructions and fixation indicators presented on the monitor’s gray background were colored so that all stimuli were isoluminant, i.e. they did not change the overall amount of light arriving at the pupil.

#### Experimental Paradigm

Participants performed 3 different tasks in separate counterbalanced blocks. In the *motor execution* task, they were instructed to reach to one of the 3 marked target positions on the table, while in the *motor imagery* task they had to *imagine* performing a reach movement to one of these targets. Finally, in the *non-motor imagery* task, they had to imagine a scene painting (*Impression, Sunrise* by Claude Monet) that had been shown to them for 30 s before the beginning of the respective block.

A schematic depiction of the setup, together with a timeline of each condition is shown in Figure 1. Each trial began with the instruction to press and hold down the start button on the touchpad. For the motor execution and motor imagery tasks, this button press triggered the presentation of a green number at the center of the monitor for 1 s, indicating the target position (“1”, “2”, or “3” for the near, middle and far target, respectively) or a resting trial (“0”). In the non-motor imagery task, a similar cue specified whether participants should imagine the scene (“I”) or rest (“0”) in this trial. After this cue, a green fixation cross (length of cross bars: 20 pixels) was shown in the center of the monitor, which changed its color to red after 2 s, serving as the “go” cue for the participant, and remained on the monitor until the end of the trial. Participants had to release the start button within a window of 3 s after the go-cue. In the motor execution task, they then reached to the specified target, returned to the start button and pressed it down. In the motor imagery task, participants imagined performing the reach movement to the specified target position and back to the start button, and pressed the start button down again when their imagined return movement had ended. The second button press allowed us to compute executed and imagined movement durations, including the hand movement back to the start position. To ensure that participants knew the target positions and established a sensorimotor representation of the reaching movement, they executed one practice reaching trial to each of the 3 targets just before the beginning of each motor execution and motor imagery task. In the non-motor imagery task, participants simply imagined the previously shown painting after releasing the start button, and they were not asked to press it down again at the end of the trial.

**Figure 1.**
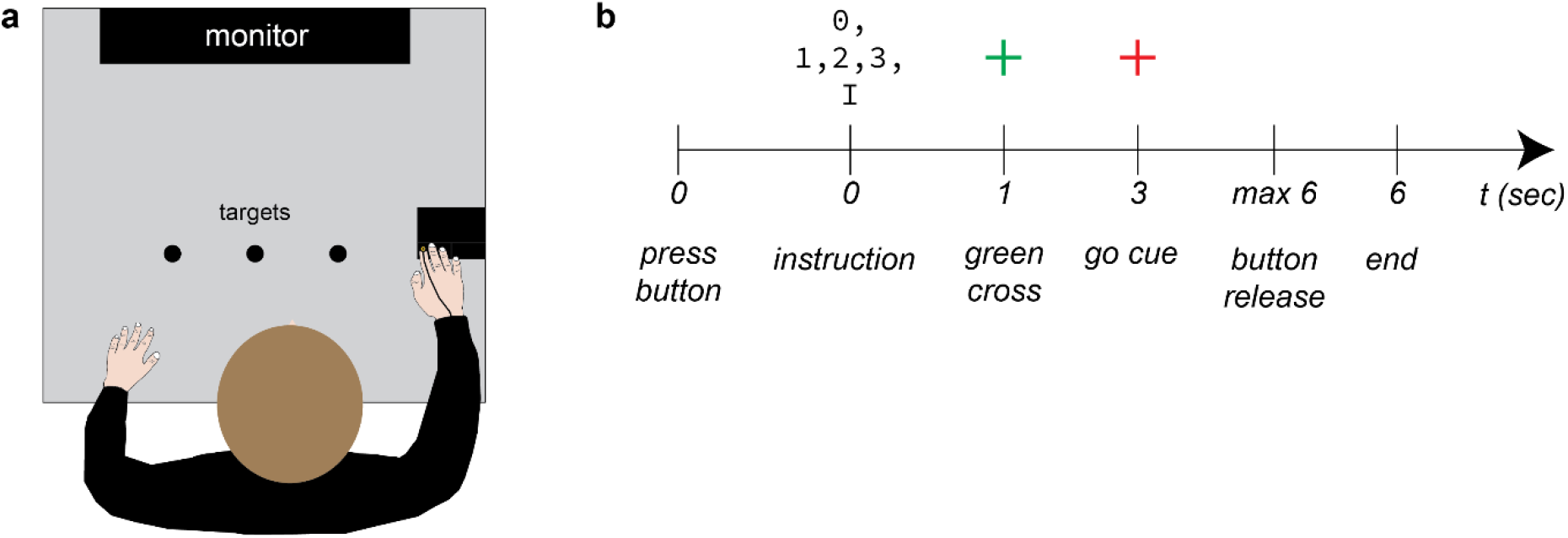
Experimental setup and timeline. (a) Top view of the setup, including the start and 3 target positions as well as the motion tracking marker on the participant’s right index finger. (b) Timeline of a single trial.

If a participant failed to release the start button within the 3 s window, the trial was canceled and repeated immediately. If a resting trial was indicated, participants were to release the start button but then do nothing specific, resting their index finger on the released button. Participants were instructed to have their eyes open during the experiment and to keep their gaze on the fixation cross for a period of 6 s after the onset of the go-cue (i.e., while the cross was red) during which pupil size was recorded. Each participant performed 60 trials each in the motor execution and motor imagery tasks (15 repetitions of each target distance, plus 15 resting trials) and 30 trials in the non-motor imagery task (15 repetitions each of imagery and rest trials). The entire experiment took approximately 45 minutes.

### Data Processing and Analysis

#### Movement times

We first examined the modulation of movement times by target distance during the motor execution and motor imagery tasks. To this end, we determined the onset of the (executed and imaginary) movement in each trial as the moment when participants released the start button following the go-cue. The end of the movement was based on when the start button was pressed *after* movement onset. The temporal difference between movement end and movement onset reflects (executed and imaginary) movement times, and this measure considers both the movement toward the target and the returning movement back to the start button. Movement times were first calculated for each trial, and then averaged across trials, separately for each target distance, task and participant. The movement time data of one participant were not correctly recorded due to a problem with the start button. Therefore, all analyses on executed and imaginary movement times are based on the remaining 29 valid datasets.

To examine possible effects of motor execution and motor imagery, as well as of target distance on movement times, we conducted a 2 (task) x 3 (distance) repeated measures ANOVA in JASP (version 0.14.1.0). Main effects of distance and/or possible interactions were followed-up with suitable post-hoc tests (one-way ANOVA and/or t-tests), corrected for multiple comparisons using the Bonferroni-Holm procedure. Effect sizes are reported as eta-squared (for ANOVAs) and Cohen’s d (t-tests). To evaluate the relationship between executed and imagined movement times, we calculated Pearson’s correlation coefficients between participants’ average movement times in both tasks (one averaged value for each target distance) using the function *corrcoef* in MATLAB 2020b (MathWorks, Natick, MA, USA).

#### Pupillary response pre-processing

Before analyzing the pupillary response characteristics and their temporal evolution, we pre-processed the pupillary data. For each participant and block of trials, we initially obtained a single Eyelink II EDF file with pupil size measures, which we processed in MATLAB R2020b. The conversion of this EDF file was done with the Edf2Mat Matlab Toolbox designed and developed by Adrian Etter at the University of Zurich. We then removed all data points 50 ms before and 50 ms after the onset and end of each blink as detected by the manufacturer’s algorithm. All missing data were then interpolated using a piecewise cubic hermite interpolating polynomial and then the data was passed through a low pass filter at 4 Hz. To allow comparison of pupil responses across trials, participants and tasks, we normalized the pupil time course of each block to zero mean and unit variance (z-score). We then segregated each file to different epochs, one per trial. Each of these epochs started 250 ms before the moment of start button release and lasted for 5000 ms after the moment of button release. Each resulting time course was then normalized to the average z-scored pupil size during the last 250 ms prior to the onset of the button release (subtractive baseline correction; see Mathôt et al., 2018).

In a second step, we used the trial-wise data to calculate the average pupillary response across the 15 repetitions of each condition that was presented within each block. This yielded a total of 10 pupillary time courses per participant: four time courses for each of the motor execution and motor imagery tasks (near, middle, far targets, and resting condition) plus two time courses for the non-motor imagery task (painting and resting condition). We then normalized the time courses of the experimental conditions by subtracting the average resting pupillary response obtained from the corresponding block of trials. Therefore, for each participant, we obtained three normalized pupillary response time courses for each motor execution and motor imagery task (one per target distance), and another one for the non-motor imagery task. These seven normalized time courses per participant reflect possible effects of task and, where applicable, target distance in the pupillary response, independent of constant factors such as random variations during resting trials, the go-cue and the button release.

#### Pupillary response characteristics

To examine whether pupillary response characteristics are affected by motor execution and motor imagery, as well as by their underlying task dynamics, we determined two main variables: the *peak response* was calculated as the largest value of each normalized time course occurring after movement onset, and the *peak response latency* was calculated as the moment relative to start button release when the peak response occurred. These values were computed separately for each target distance, task and participant based on the averaged time courses across the trials of each condition because the noisy individual recordings did not allow to identify a clear peak on each trial separately.

We first tested whether motor execution and motor imagery caused a change in pupil diameter relative to resting trials. To this end, we used one-sided one-sample t-tests to contrast the peak response in each of the six conditions against zero (i.e. average baseline pupillary response). We used one-sided t-tests because we hypothesized pupil dilations compared to baseline during motor execution (e.g., Moresi et al., 2008a; Rozado et al., 2017), during motor-imagery (e.g., Rozado et al., 2015; Rozado et al., 2017) and during cognitive tasks (e.g., Einhauser, 2017; Hess & Polt, 1964). In a second step, possible effects of task and target distance on each of these variables were examined with separate 2 (task) x 3 (distance) ANOVAs, using suitable post-hoc testing as described earlier.

#### Pupillary response temporal evolution

We further examined the temporal evolution of the pupillary responses between the three target distances in the motor imagery and motor execution tasks, using sample - wise analysis. First, we tested whether the time courses of the pupillary responses during the three main tasks were different from those during the resting baseline, and if so, at which times during the trial. To this end, we use one-sided one-sample t-tests to compare time courses against zero at each sample time point (2 ms interval at 500 Hz), where zero represents the average baseline pupillary response after subtraction of the baseline time course. Second, to investigate possible differences in the temporal pupil dynamics between motor execution and motor imagery across target distances, as well as interactions between these two factors, we ran linear mixed models (LMM) separately for each sample time point in R (version 4.2.1) using the *lme4* (version 1.1-31) and *lmerTest* packages (version 3.1-3). Associated post-hoc pairwise comparisons were performed using the *emmeans* package (version 1.8.2), applying Satterthwaite’s method to estimate degrees of freedom (df) and correcting for multiple comparisons within each time point model using Bonferroni-Holm correction. LMMs modeled fixed effects of task and target distance as well as their interaction, and random intercepts were included for each participant. LMMs here allowed to optimally model trial-by-trial variability in the pupil time course at each time step and therefore we included all individual trials, instead of the average responses for each condition. Due to the large number of samples and comparisons for the time course analyses, we controlled the False Positive Rate at the 5% level for both LMM and t-test results (false discovery rate or FDR correction; Benjamini & Hochberg, 1995).

#### Pupillary response across motor and non-motor tasks

It is possible that any pupillary responses found during motor imagery are related to non-motor, cognitive factors, such as memory operations (e.g., Naber et al., 2013; Kucewicz et al., 2018). To assess this, we tested whether the pupillary peak response as well as the average pupillary time course differed between the three main tasks (motor execution, motor imagery, non-motor imagery). To this end, we first calculated a single normalized time course separately for each task and participant, averaging across the 3 target distances (45 trials) for the motor execution and motor imagery tasks. For the non-motor imagery task, we averaged across the 15 trials involving imagery of the previously shown scene. We then calculated the peak response after movement onset for each task and used three one-sided one-sample t-tests to examine whether this was different from zero (i.e. from the average baseline pupillary response). Task effects on the peak pupillary response were examined with a LMM that had fixed effects of task and random effects of participant, while main effects of task were explored with Bonferroni-Holm corrected post-hoc t-tests.

We further compared each task’s average pupillary response profile to zero using one-sided one-sample t-tests for each sample, as described above, to determine which tasks produced a pupil dilation different from rest, and for which time periods. We also conducted sample-by-sample LMMs with a fixed effect of task and random intercepts per participant in order to identify possible periods during which there were differences between tasks. Possible effects were further explored with post-hoc t-tests that were FDR-corrected for multiple comparisons, as described earlier.

## Results

### Effects of motor execution and motor imagery

#### Movement times

We first examined whether executed and imaginary movement times were modulated by target distance. Movement times were generally longer during motor imagery than during motor execution (F_1, 28_ = 8.96, p = 0.006, η^2^ = 0.09; Figure 2a). Unsurprisingly, they were also affected by target distance (F_2, 56_ = 109.73, p < 0.001, η^2^ = 0.39): they were shorter for near than middle (t_28_ = −8.63, p < 0.001, d = −1.61) and far (t_28_ = − 14.74, p < 0.001, d = −2.74) targets, as well as for middle than far targets (t_28_ = −6.11, p < 0.001, d = −1.12), in line with previous work showing modulation of movement time by movement distance (e.g., Voudouris et al., 2010). Importantly, there was no interaction between task and target distance (F_2, 56_ = 2.31, p = 0.108, η^2^ = 0.01), suggesting that movement time modulation by target distance was similar for both motor execution and motor imagery tasks.

**Figure 2.**
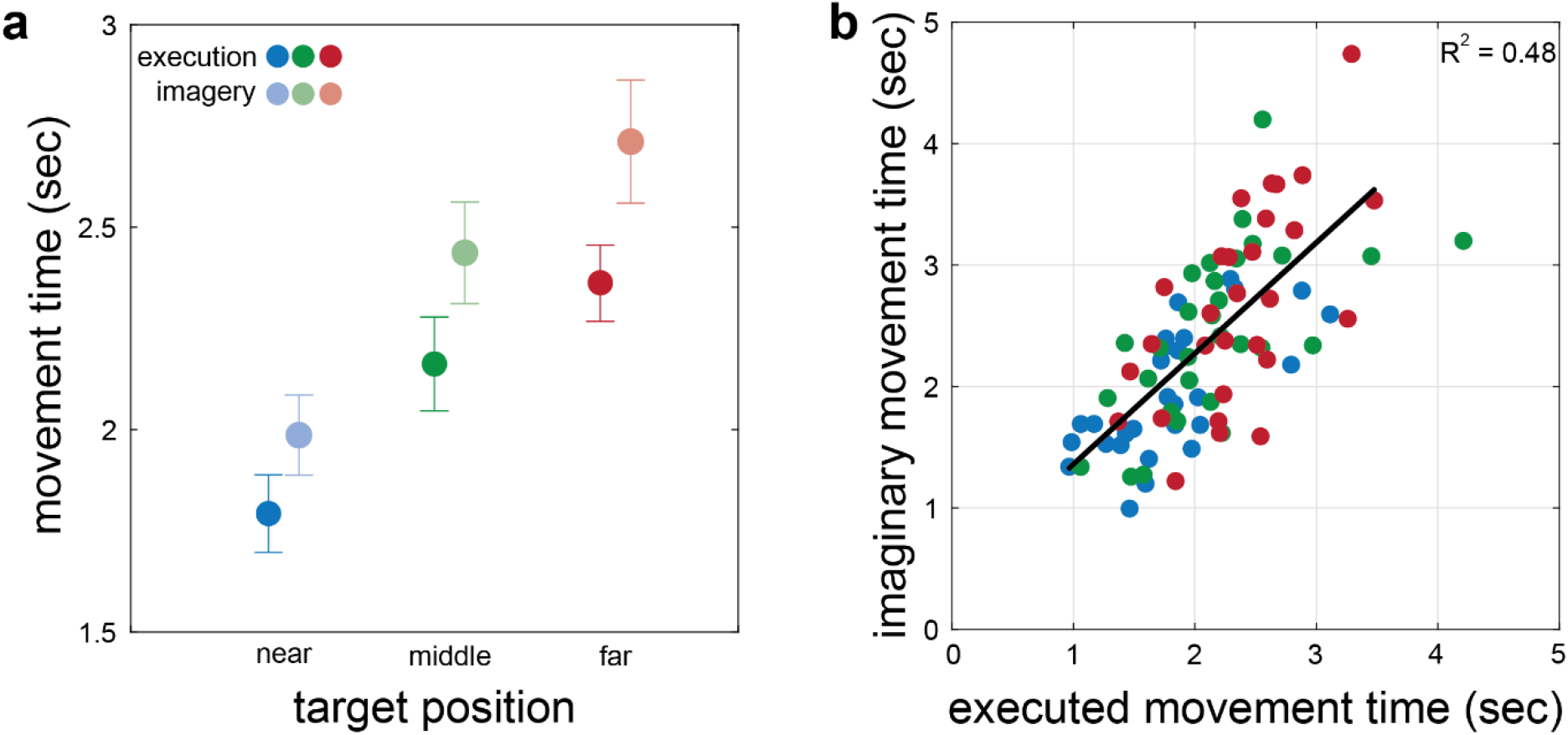
Movement time. (a) Effects of motor task and target distance on executed and imagined movement times. Averages and standard errors of the mean are shown. (b) Relationship between executed and imagined movement times. Each symbol in (b) shows the average movement times of individual participants, color-coded for target distances. Regression line demonstrates the relationship between the movement times.

There was also a strong correlation between executed and imaginary movement times (r = 0.69, p < 0.001, R² = .48; Figure 2b), in line with previous work (e.g., Papaxanthis et al., 2002; Munzert et al., 2015). This confirms that participants were engaged in performing the motor imagery task. After this sanity check, we examined whether pupillary responses followed a similar modulation by the task and the target distance.

#### Peak pupillary responses

We then tested whether motor execution and motor imagery led to a change in pupil dilation relative to the resting condition. To this end, we contrasted the peak pupillary response for each motor task and target distance against zero (i.e. respective resting activity). Pupil dilations were evident during motor execution for all 3 target distances (near: t_29_ = 13.01, p < 0.001, d = 2.37; middle: t_29_ = 15.34, p < 0.001, d = 2.81; far: t_29_ = 15.16, p < 0.001, d = 2.79; Figure 3a). Similarly, motor imagery led to pupil dilation irrespective of target distance (near: t_29_ = 9.38, p < 0.001, d = 1.71; middle: t_29_ = 8.66, p < 0.001, d = 1.58; far: t_29_ = 7.71, p < 0.001, d = 1.41; Figure 3a).

**Figure 3.**
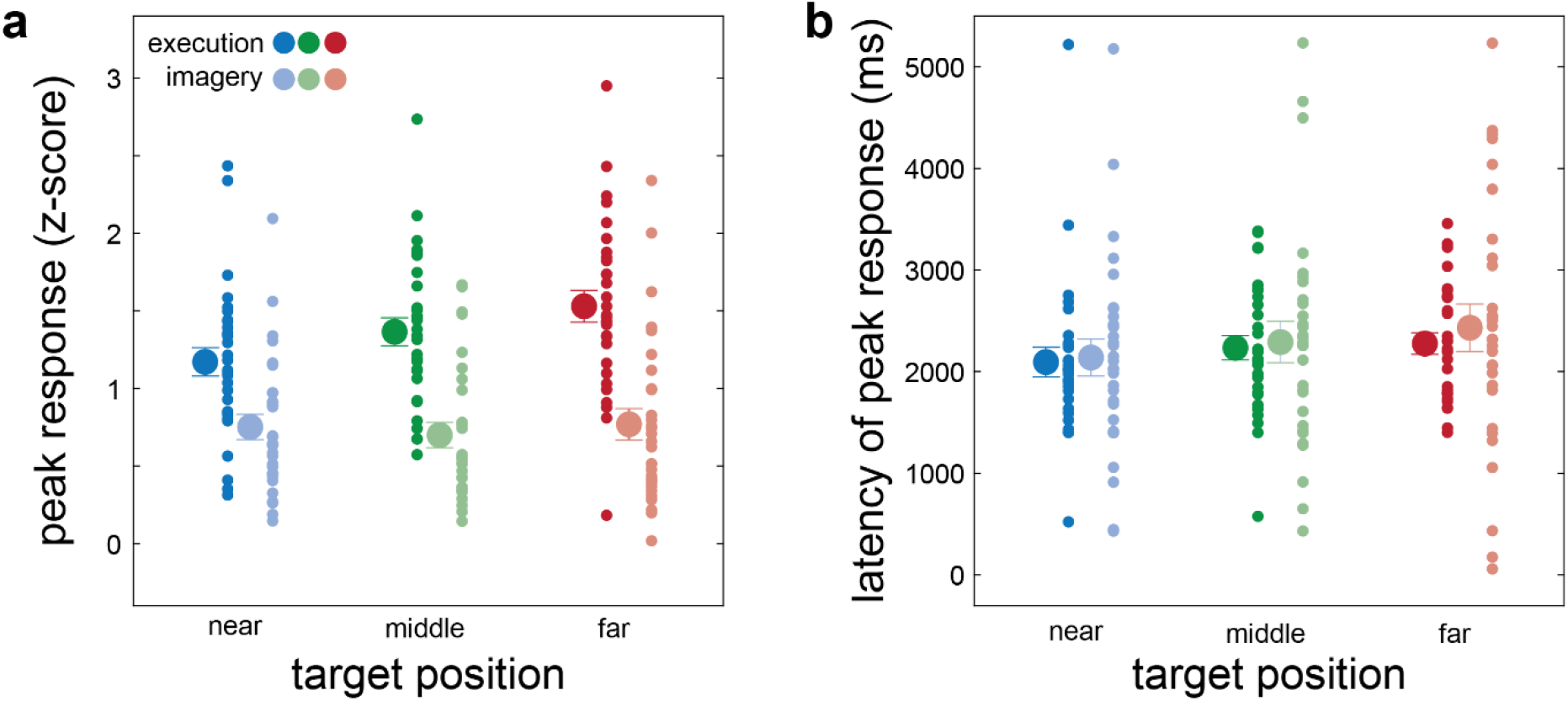
Response characteristics. Effects of motor execution and motor imagery, as well as of target distance on executed and imagined (a) peak pupillary response, and (b) peak response latency. Averages and standard errors of the mean are shown, with individual participants depicted with smaller symbols.

We then examined whether peak pupillary responses, as well as their latency, were different between motor execution and motor imagery, and between the 3 target distances. The *peak pupillary response* was greater during motor execution than motor imagery (F_1, 29_ = 52.55, p < 0.001, η^2^ = 0.51; Figure 3a). It was also affected by target distance (F_2, 58_ = 14.72, p < 0.001, η^2^ = 0.03), but we will interpret this effect within the significant interaction between task and target distance (F_2, 58_ = 9.74, p < 0.001, η^2^ = 0.03). This interaction showed that the peak pupillary response was modulated by the target’s distance only during motor execution (F_2, 58_ = 19.08, p < 0.001, η^2^ = 0.39), but not during motor imagery (F_2, 58_ = 1.16, p = 0.322, η^2^ = 0.38). Specifically, during motor execution, peak pupillary responses were scaled to target distance, as they were larger for middle than near (t_29_ = 3.34, p = 0.004, d = 0.61), far than near (t_29_ = 6.17, p < 0.001, d = 1.12) and far than middle targets (t_29_ = 2.83, p = 0.019, d = 2.83).

The *peak response latency* was similar between motor execution and motor imagery (F_1, 29_ = 0.19, p = 0.665, η^2^ = 0.01; Figure 3b), as well as across the 3 target distances (F_2, 58_ = 1.99, p = 0.145, η^2^ = 0.02). In addition, there was no interaction between these 2 factors (F_2, 58_ = 0.16, p = 0.848, η^2^ = 0.01).

In short, both motor execution and motor imagery led to significant pupil dilations compared to rest, but the timing of these peak dilations remained invariant across all conditions. Pupillary dilations were greater when executing a movement rather than just imagining it. In addition to the finding that peak dilations increased with longer executed but not with longer imagined movements, these findings suggest that pupil responses are sensitive to changes in the musculoskeletal system, for instance in muscle activation or proprioceptive and/or tactile feedback associated with the executed movement.

#### Temporal evolution of pupillary responses

We also examined the temporal evolution of the pupillary responses associated with motor execution and motor imagery. Specifically, we were interested in whether the average time series of pupillary dilation are affected by the underlying task and its dynamics (i.e. target distance). This time series analysis expands on the peak dilation metrics in the previous section and allows to characterize the temporal dynamics of pupillary responses, for instance by determining the precise on- and offsets of motor imagery-related pupil dilation. These time ranges could further serve as markers of whether an imagery task was performed within the correct time interval in future studies.

We first tested whether pupillary responses during all motor execution and motor imagery conditions were different from the associated resting baseline. To this end, we first conducted sample-by-sample t-tests against zero throughout each time course, which revealed that all 6 responses differed from the resting baseline for substantial durations (Figure 4a). In detail, executed movements caused significant dilations that were evident already at 118, 92, and 58 ms for the near, middle and far target, respectively, and that remained systematically uninterrupted until 3574, 5000, and 5000 ms relative to the start button release (all t > 1.72, all p < 0.049). Note that pupil time series were analyzed until 5000 ms after the start button release, therefore responses when reaching to the middle and far targets likely lasted even longer. Meanwhile, motor imagery also led to significant dilations that were evident at 166, 186, and 194 ms and that lasted until 3038, 3096, and 3422 ms for the near, middle and far target location, respectively (all t > 1.94, all p < 0.049). Thus, both tasks led to systematic pupillary responses, which lasted longer in the motor execution than motor imagery task. They also lasted longer for targets farther away. This is in line with the previously reported finding of larger peak pupillary responses during motor execution and for greater target distances (Figure 3b).

**Figure 4.**
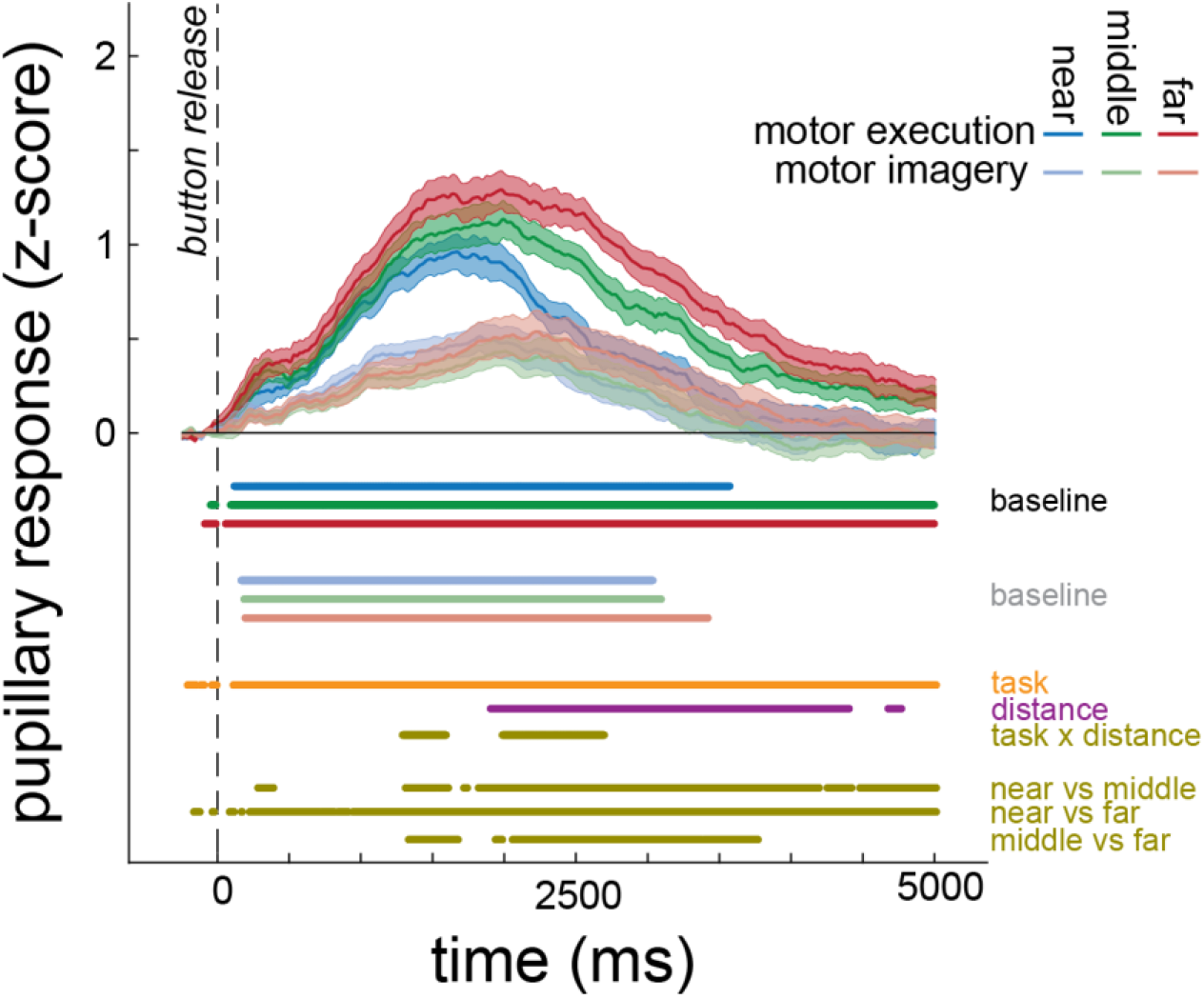
Time course of pupil dilation response. Normalized time courses (after z-scoring and subtraction of resting pupil time course) for motor execution and motor imagery, separately for each of the 3 target distances. The first 6 horizontal bars illustrate the moments when the responses are significantly greater than the resting baseline, color-coded by target distance. Pupillary responses during both tasks increase relative to the resting baseline (black horizontal line), for all target distances. Main effects of task and distance, as well as interactions are shown with the subsequent 3 horizontal bars. The effect of distance is interpreted within the interaction, which revealed that the effect of distance was evident only during motor execution. Associated post-hoc t-tests across the 3 distances during motor execution are shown with the last 3 horizontal bars. Each trace is the average response along with the standard error of the mean. No post-hoc test bars are shown for the motor imagery condition, as no differences were found. Vertical dashed line indicates start button release.

In a next step, we explicitly tested when the time courses of different tasks (motor execution and imagery) and target distances (near, middle, far) differed significantly, as well as for any interaction between them. Pupillary responses were systematically stronger during motor execution than motor imagery between 108 and 5000 ms after movement onset (all F > 4.08, all p < 0.047), though systematic differences were evident also earlier, and even before movement onset. In addition, systematic differences across the target distances were evident between 1898 until 4394 ms relative to movement onset (all F > 3.81, all p < 0.049). Because there was also an interaction between task and target distance between 1286 and 1590 ms, as well as between 1980 and 2690 ms after movement onset, we interpret the main effect of distance within the interaction effect by running separate pairwise t-tests between target distances, separately for each task. For the motor execution task, there was a clear effect of target distance with farther away targets leading to stronger pupil dilations. More specifically, pupil dilations were larger when reaching to the middle compared to the near target, and this was generally evident between 1306 and 5000 ms after movement onset (all t > 0.39, all p < 0.049). Similarly, pupil dilations were larger when reaching to the far than to the near target, already starting 224 ms after movement onset and until the end of the recording (all t > 0.66, all p < 0.049). Pupil responses were stronger also during reaching to the far as compared to the middle target, particularly between 1938 ms after movement onset and until the end of the recording (all t > 0.29, all p < 0.049). For the motor imagery task, however, there was no single moment when the time series significantly differed between pairs of target distances (all t < 0.12, all p > 0.99), in line with the results reported in the previous section about the peak pupillary dilation.

In sum, the time course analyses revealed that pupillary dilations are larger during both motor execution and motor imagery when compared to rest. Pupillary responses were generally stronger and lasted longer for farther away targets, however the interaction demonstrates that this was the case only during movement execution and not during motor imagery.

### Comparing across tasks

The pupillary dilations during motor imagery may also reflect other cognitive processes independent of the internally simulated movement dynamics. To investigate this possibility, we included a non-motor imagery task in our experimental design, where participants had to imagine the details of a previously shown painting. We compared average pupillary responses across the 3 tasks (motor execution, motor imagery, non-motor imagery). To this end, we calculated the average response time course in each of the 3 tasks (motor execution, motor imagery, and scene imagery) for each participant, averaging across target distances in the motor execution and motor imagery tasks. We then calculated and compared the peak dilation response for each task. As expected from the previous results, the peak pupillary response was greater than the resting baseline during motor execution (t_29_ = 14.72, p < 0.001, d = 2.68) and motor imagery (t_29_ = 8.82, p < 0.001, d = 1.61; Figure 5a). We also found that the peak pupillary response was greater than the resting baseline during the non-motor imagery task (t_29_ = 6.53, p < 0.001, d = 1.19; Figure 5a). Peak pupillary dilation was also different across tasks (F_2, 58_ = 28.48, p < 0.001, η^2^ = 0.49): it was larger during motor execution than motor imagery (t_29_ = 5.73, p < 0.001, d = 1.04) and non-motor imagery (t_29_ = 7.12, p < 0.001, d = 1.30). Peak dilation was similar between motor imagery and non-motor imagery (t_29_ = 1.39, p = 0.170, d = 0.254).

**Figure 5.**
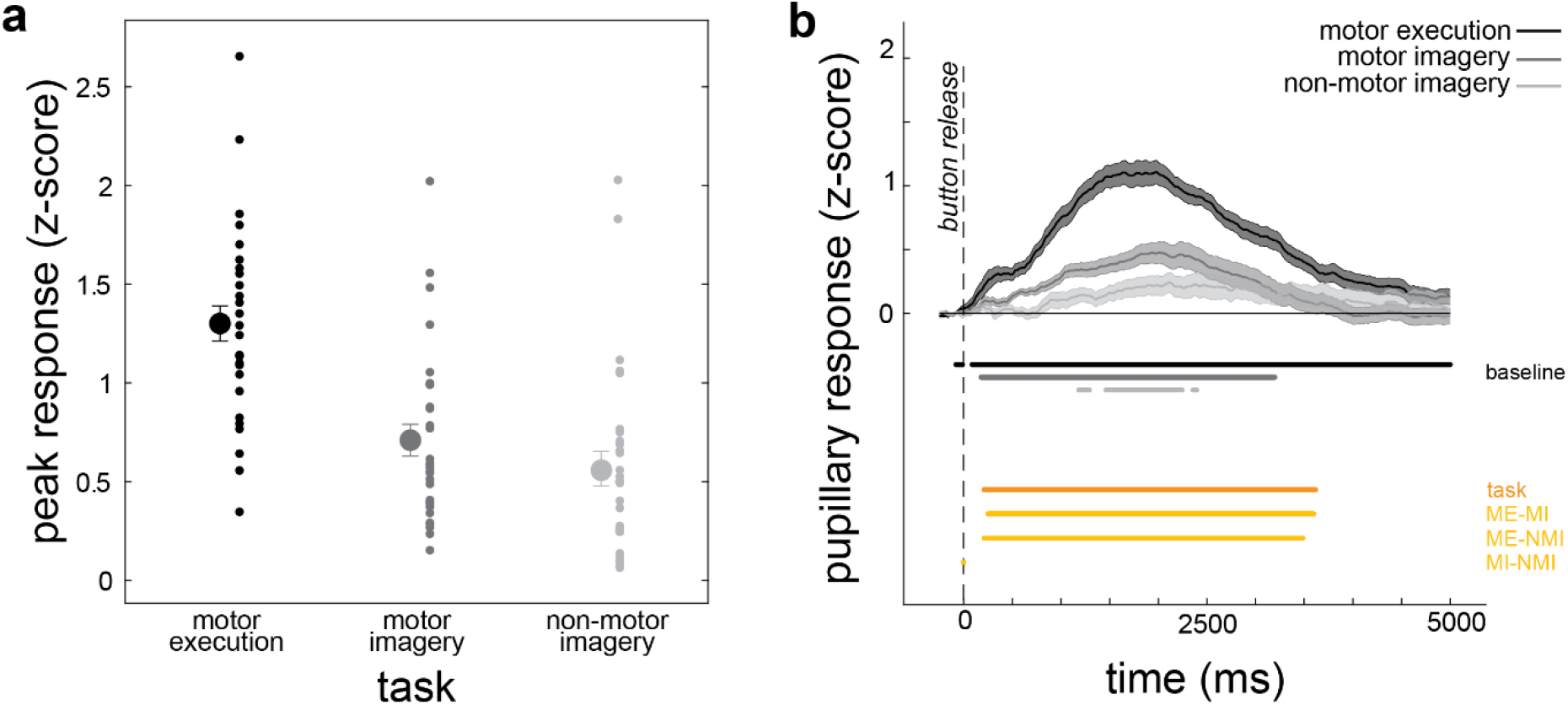
Peak pupil dilations and temporal evolution across tasks. (a) Peak pupillary response and (b) normalized time courses for motor execution, motor imagery and non-motor imagery tasks. The LMM analysis on the time courses revealed a main effect of task (red line) that is explored with post-hoc t-tests (ME: motor execution; MI: motor imagery; NMI: non-motor imagery). Details as in Figure 4.

We also examined the time course of the pupillary responses associated with the 3 main tasks (Figure 5b). First, we tested whether the time courses were different from baseline, and if so at which time periods. The motor execution task led to dilations starting already 80 ms before movement onset and remaining systematic until the end of the recording (all t > 1.73, all p < 0.049; Figure 5a). The motor imagery task also led to dilations but for a shorter period of time, specifically between 176 and 3198 ms after movement onset (all t > 1.98, all p < 0.049). Significant dilations were also evident during the non-motor imagery task, but these were of an even shorter duration, namely between 1190 and 2394 ms after movement onset (all t > 2.51, all p < 0.049). We also investigated whether the time courses were different between the 3 tasks. The LMM analysis revealed that there was a main effect of task between 224 and 3616 ms after movement onset (all F > 3.58, all p < 0.049; Figure 5b): Responses during motor execution were greater than those during the motor imagery task (248 and 3598 ms after movement onset; all t > 0.048, all p < 0.048) and the non-motor imagery task (218 and 3486 ms after movement onset; all t > 0.048, all p < 0.048). However, there was no systematic difference between the motor and non-motor imagery tasks (all t < 0.001, all p > 0.99). These are in line with the effects on the peak pupillary response.

## Discussion

In Experiment 1, we show that both execution and imagery of a goal-directed reach movement led to significant pupil dilation compared to rest. Dilations during motor execution started already ∼100 ms before movement onset and lasted for at least 5000 ms, whereas dilation during motor imagery began ∼180 ms after the onset of imaginary movement and lasted for ∼3000 ms. In addition, pupillary dilations were larger for farther away targets, but this was the case only when executing the movement. Previous work has shown that pupillary responses are stronger when performing more complex movements or when applying more force (Richer & Beatty, 1985; Zenon et al., 2014) and thus when muscular activation is stronger. This could explain not only the stronger pupillary responses with longer movement distances, but also the stronger responses during motor execution compared to motor imagery, as muscular activations during motor execution were likely stronger for farther away targets and relative to motor imagery. As we did not measure muscle activity in Experiment 1, this explanation remains speculative.

The second main finding of Experiment 1 is that pupillary responses were similar between motor imagery and non-motor imagery tasks, as reflected both in the peak of the response (Figure 5a) and in the time courses (Figure 5b). It is known that non-motor cognitive tasks such as problem-solving, mental multiplication and mental rotation can cause pupil dilations (e.g., Brauer et al., 2021; Einhauser, 2017; Hess & Polt, 1964; Schuetz et al., 2021). Imagining the previously shown painting during the non-motor imagery task entails memory processes that likely also lead to additional pupil dilations. Considering the similarity in the responses between motor and non-motor imagery tasks in our experiment, we cannot exclude the possibility that pupil dilations during the motor imagery task reflect only or partly cognitive-related process of the previously executed reaching movements, such as recalling the target position or movement duration from memory.

A difference between the motor and non-motor tasks was that participants had to press a button at the end of the motor tasks, but not at the end of the non-motor imagery task. This additional button press was necessary to evaluate actual and imagined movement times and to relate our results to traditional findings from mental chronometry paradigms (Decety & Jeannerod, 1996; Papaxanthis et al., 2002), and to confirm that participants engage in motor imagery, irrespectively of their pupillary responses. Yet, the button press at the end of each trial may have caused an anticipatory elevation of the pupillary response (Richer & Beatty, 1985; Hupé et al, 2009), which could (at least partially) explain the dilation found in the motor imagery condition. To rule out this explanation, we conducted a second experiment that did not include a button press at the end of a trial. If pupil dilations stem from processes other than the anticipation of the upcoming button press, then we again expect to find pronounced dilations during both motor execution and motor imagery in Experiment 2.

## Experiment 2

### Methods

In Experiment 2, we examined whether the pupillary responses during motor execution and motor imagery obtained in Experiment 1 may reflect an anticipatory dilation associated with the upcoming button press at the end of each trial (Richer & Beatty, 1985; Hupé et al, 2009). Additionally, we investigated whether any possible pupillary responses during motor imagery might have been caused by muscular activity associated with the underlying imagined movement. To this end, we asked a new set of participants to perform only the motor execution and motor imagery tasks toward the same 3 targets, identical to those presented in Experiment 1, but now without any button press at the end of each trial. In addition, we recorded muscular activity from their arm that was involved in the task. If pupillary dilations during motor execution and motor imagery were not evoked primarily by the anticipated button press, then they should again be different from the resting baseline pupillary time course in both tasks. In addition, if pupillary responses are related to muscular activity produced by the effector involved, even in the absence of a discrete movement, then the strength of the pupillary response should relate to the strength of the muscular activation. The design, apparatus, data and statistical analyses for Experiment 2 were identical to those of Experiment 1, except the aspects described below.

### Participants

For this experiment, we recruited a sample of 30 participants (19 women, 11 men, age range: 18-39 years), all right-handed according to the German translation of the Edinburgh Handedness Inventory (range: 50-100; Oldfield, 1971). None of the participants previously participated in Experiment 1.

### Experimental Setup

In addition to the measures recorded in Experiment 1, we also recorded electromyographic (EMG) activity from the right upper arm. An eight-channel amplifier (actiCHamp, BrainProducts GmbH, Gilching, Germany) in combination with the software PyCorder (ver. 1.0.9) was used to record EMG activity from the participants’ right anterior deltoid at 2000 Hz. We recorded from the deltoid muscle because it shows high sensitivity in detecting upper-arm movements (e.g., Trigili et al., 2019) and because its activity scales well with reaching distance (e.g., Buneo et al., 1994). Two bipolar electrodes were placed at the intermediate part of the belly of the right deltoid with an interelectrode distance of 1 cm. A single ground electrode was fixed to the ulnar styloid process of the left hand that was always resting on the table.

### Experimental Paradigm

The instruction of Experiment 2 was changed by asking participants to “return to the button *without* pressing it”. The start button could not be pressed down unless intentionally applying downward force on it. Thus, there were no special requirements for participants to avoid pressing the button upon finishing each trial.

### Data Processing and Analysis

EMG data analysis were performed as follows: For each participant, we obtained a single file with muscular activity measures for each task. Each of these files was first band-pass filtered with cut-offs at 20 and 900 Hz, and then full-wave rectified. We then segregated this file into different epochs, one for each trial. Each of these epochs started 500 ms before the start button release, and lasted for 5000 ms after the moment of start button release. Each trial was first normalized by subtracting the average EMG activity during the last 500 ms prior to button release. We then averaged the EMG time courses across the trials, separately for each condition and participant, obtaining 8 maxima per participant: 3 for each of the motor execution and motor imagery condition, and 2 for the resting trials of the respective block. We then normalized the maximum EMG activity by subtracting the average maximal resting EMG activity from the maximal EMG activity during the motor execution and motor imagery trials of the respective block. To examine whether motor execution and motor imagery led to EMG activity relative to the resting baseline, we submitted the normalized maximal EMG values to 6 separate one-sided one-sample t-tests. Please note that we use one-sided t-tests because we have rectified the EMG signal and thus muscle activity cannot be negative. To assess effects of task and target distance on the normalized maximal EMG activity, we submitted the respective values to a 2 (task) x 3 (target distance) ANOVA, and performed suitable Bonferroni-Holm corrected post-hoc testing whenever necessary. To test whether pupillary response characteristics depend on muscular activity, we computed Pearson’s correlation coefficients between the normalized maximal EMG value and the peak pupillary response.

## Results

### Peak pupillary responses

Similar to Experiment 1, peak pupillary responses were significantly larger than the resting baseline during motor execution for all 3 target distances (near: t_29_ = 10.51, p < 0.001, d = 1.91; middle: t_29_ = 11.67, p < 0.001, d = 2.13; far: t_29_ = 14.01, p < 0.001, d = 2.56; Figure 6a). Likewise, pupil dilation was greater during motor imagery to all 3 targets relative to rest (near: t_29_ = 6.99, p < 0.001, d = 1.27; middle: t_29_ = 6.46, p < 0.001, d = 1.18; far: t_29_ = 7.08, p < 0.001, d = 1.29). These results confirm the previously reported findings of Experiment 1.

**Figure 6.**
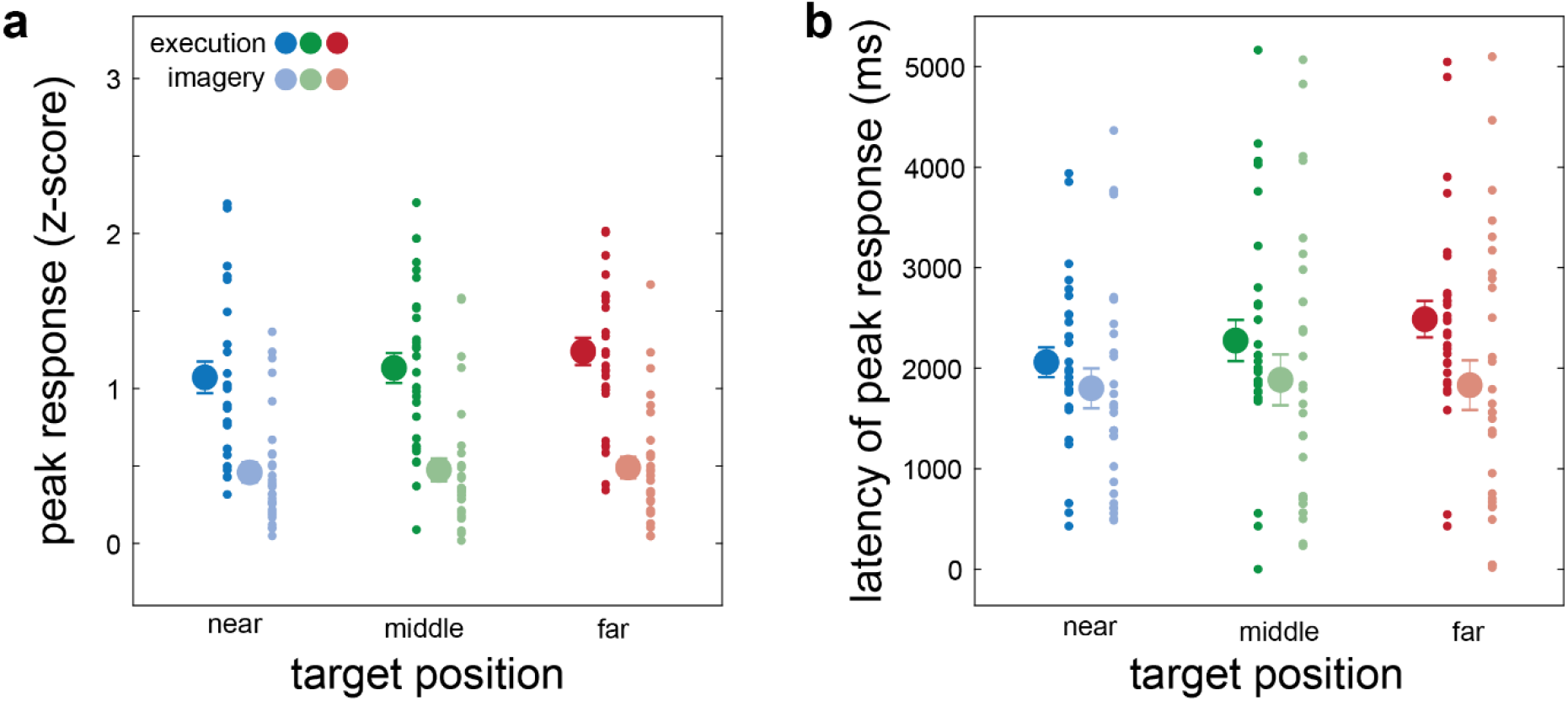
Response characteristics in Experiment 2. Effects of the 3 tasks on (a) peak pupillary response, and (b) peak response latency. Details as in Figure 3.

**Figure 7.**
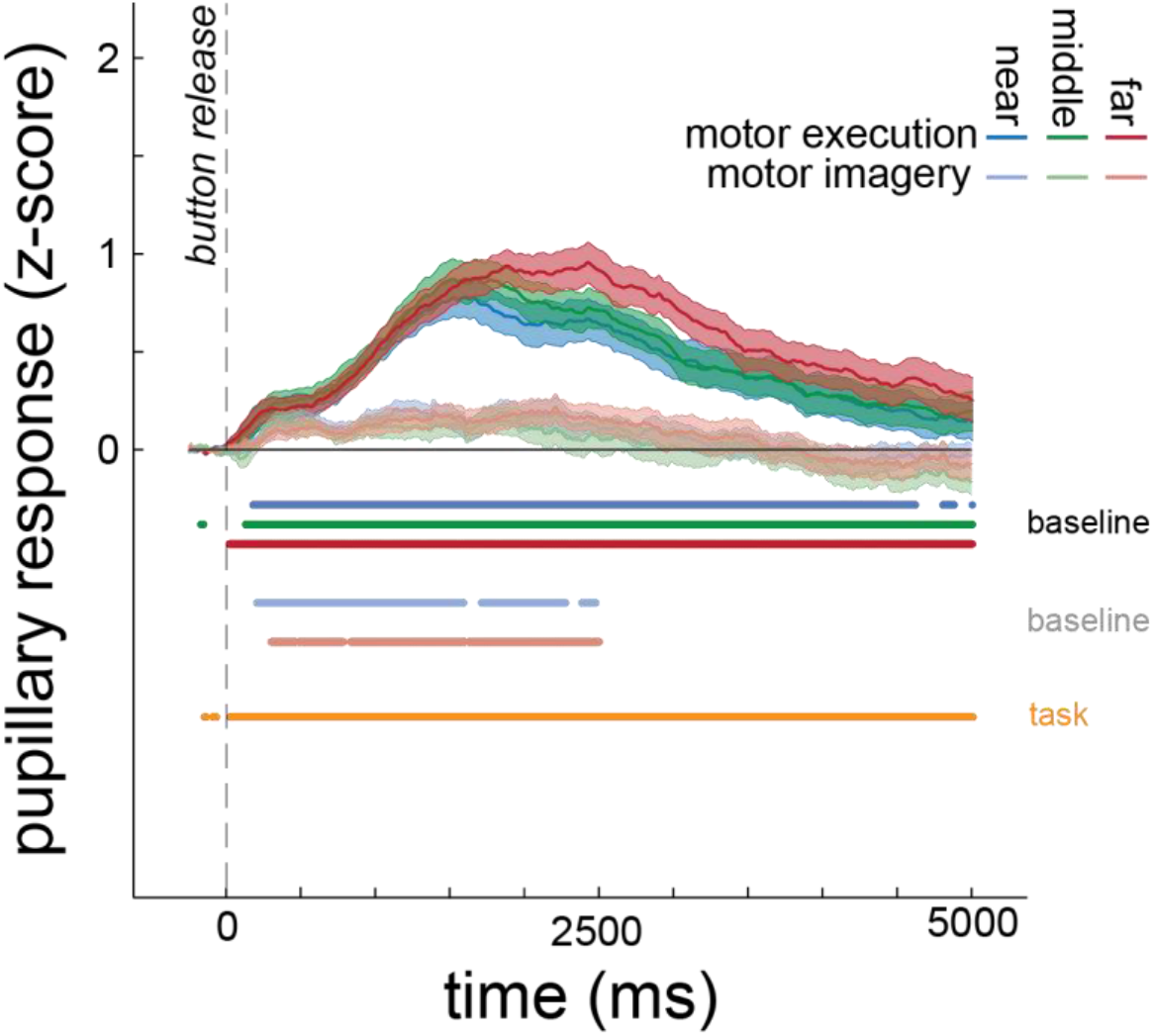
Response temporal evolution in Experiment 2. Temporal modulation of the responses for each of the 3 targets for motor execution and motor imagery. Details as in Figure 4.

There was also a main effect of task (F_1, 29_ = 46.78, p < 0.001, η^2^ = 0.53), as the peak response was greater during motor execution than motor imagery. We also found a main effect of target distance (F_2, 58_ = 4.08, p = 0.022, η^2^ = 0.01), because pupillary responses were larger for the far compared to the near target (t_29_ = 2.83, p = 0.019, d = 0.52), but no other comparisons were systematic (both |t| < 1.7, both p > 0.176). Although the effect of target distance appeared to be the case mainly in the motor execution condition, there was no interaction (F_2, 58_ = 1.53, p = 0.226, η^2^ = 0.01).

The peak response latency occurred slightly later during motor execution than during motor imagery (F_1, 29_ = 4.84, p = 0.036, η^2^ = 0.06; Figure 6b). There was neither an effect of target distance (F_2, 58_ = 1.72, p = 0.187, η^2^ = 0.01), nor an interaction (F_2, 58_ = 0.76, p = 0.469, η^2^ = 0.01).

### Temporal evolution of pupillary responses

Motor execution led to significant dilations relative to the resting baseline (all t > 1.73, all p < 0.049), which started at 176, 126 and 16 ms from the moment of start button release for the near, middle, and far target, respectively, and these responses were significant until the end of the recording. For the motor imagery task, there were again significant dilations but these only differed significantly from baseline when imagining reaches to the near and to the far target (all t > 2.14, all p < 0.049 for the above time range). The responses during imagined reaching to the middle target were qualitatively evident but not as systematic (all t < 3.43, all p > 0.054).

We then examined possible differences in pupillary time courses between motor execution and imagery, between the 3 target distances, and a possible interaction between both factors. Pupillary responses were systematically stronger during motor execution than motor imagery for almost the complete duration of the recording, starting at 42 ms after movement onset and for the entire duration of recording (all F > 3.99, all p < 0.049). This replicates the respective findings of Experiment 1. Despite the qualitatively stronger and longer responses during reaches to the far target, as in Experiment 1, there was neither a main effect of target distance (all F < 3.69, all p > 0.099) nor an interaction (all F < 1.93, all p > 0.989), similar to the results of the peak pupillary response.

### Pupillary temporal evolution across experiments

When comparing pupil dilation time series between Experiments 1 and 2, pupil dilations appear generally greater in Experiment 1 (Figure 8a, shown after averaging across all 3 movement distances). While this might reflect differences in pupil dilation relating to the removal of the button press, our data were z-scored and normalized separately for each experiment, which limits any conclusions that can be drawn from this comparison. The differences in overall dilation amplitude shown in Figure 8a thus could simply be caused by two very different samples of participants. For a better comparison, we calculated the difference in pupillary responses between motor execution and motor imagery, across all target distances and separately for each participant and experiment. This was based on the assumption that the extra button press was present in both tasks in Experiment 1 and thus could have influenced both time series, but that the relative dilation amplitudes might be comparable between tasks The resulting difference time series are shown in Figure 8b and indicate almost no difference in response between both experiments. Therefore, while the additional button press might have caused overall larger dilation across tasks of Experiment 1, these results suggest that the pupil dilations during the motor imagery task in Experiment 1 were not simply caused by the upcoming button press.

**Figure 8.**
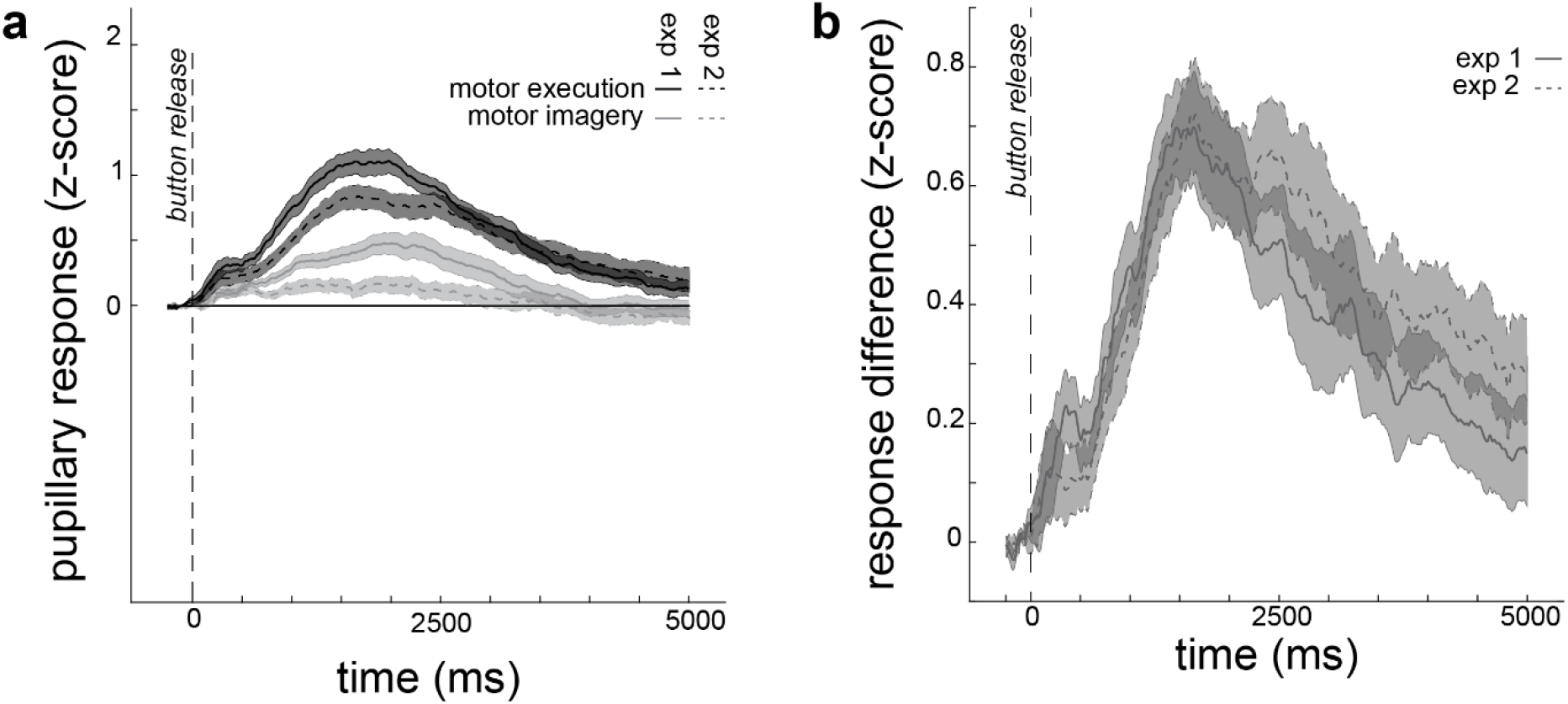
Pupillary responses between tasks and experiments. (a) Pupillary responses during motor execution and motor imagery in both experiments. (b) Difference in pupillary responses between motor execution and motor imagery, separately for each experiment.

In sum, in Experiment 2 we found systematic pupillary dilations during motor execution and motor imagery relative to baseline. This effect was present even when participants were instructed not to press the button at the end of their reach. Thus, the anticipation of the final button press cannot fully explain the results of Experiment 1, suggesting that these dilations do not simply result from an anticipated manual action.

### Muscle activity during motor execution and motor imagery

Experiment 2 also tested whether pupillary responses during motor imagery may have been related to muscle activations. We first tested whether the normalized maximal muscle activation was larger in the motor execution and imagery tasks than the muscular activity obtained during baseline (resting). To this end, we conducted 6 separate one-sided t-tests against zero, one per condition. Muscle activity was pronounced during motor execution, for all 3 target distances (near: t_29_ = 3.35, p = 0.002, d = 0.61; middle: t_29_ = 3.34, p = 0.002, d = 0.61; far: t_29_ = 3.36, p = 0.002, d = 0.61; Figure 9). As expected, EMG activity during motor imagery was not different from zero for any target distance (near: t_29_ = −0.82, p = 0.417, d = −0.15; middle: t_29_ = −0.24, p = 0.806, d = −0.04; far: t_29_ = −1.31, p = 0.200, d = −0.24; Figure 9). These findings are supported by a main effect of task on maximal normalized muscle activity, which was stronger during motor execution than motor imagery (F_1, 29_ = 14.39, p < 0.001, η^2^ = 0.24). Muscle activity was also affected by target distance (F_2, 58_ = 6.71, p = 0.002, η^2^ = 0.02), as it was larger for far than near targets (t_29_ = 3.65, p = 0.002, d = 0.66) but no other comparison was systematic (both t < 2.01, both p > 0.009, both d < 0.37). There was also a significant interaction (F_2, 58_ = 6.44, p = 0.003, η^2^ = 0.02). Post-hoc one-way ANOVAs separately for each task revealed that the effect of target distance was evident only during the motor execution (F_2, 58_ = 6.88, p = 0.002, η^2^ = 0.19) but not during motor imagery (F_2, 58_ = 0.55, p = 0.578, η^2^ = 0.02). In sum, muscular activity was negligible during motor imagery, suggesting that the effects of motor imagery on pupil diameter are unlikely to arise from underlying activations of the musculoskeletal system.

**Figure 9.**
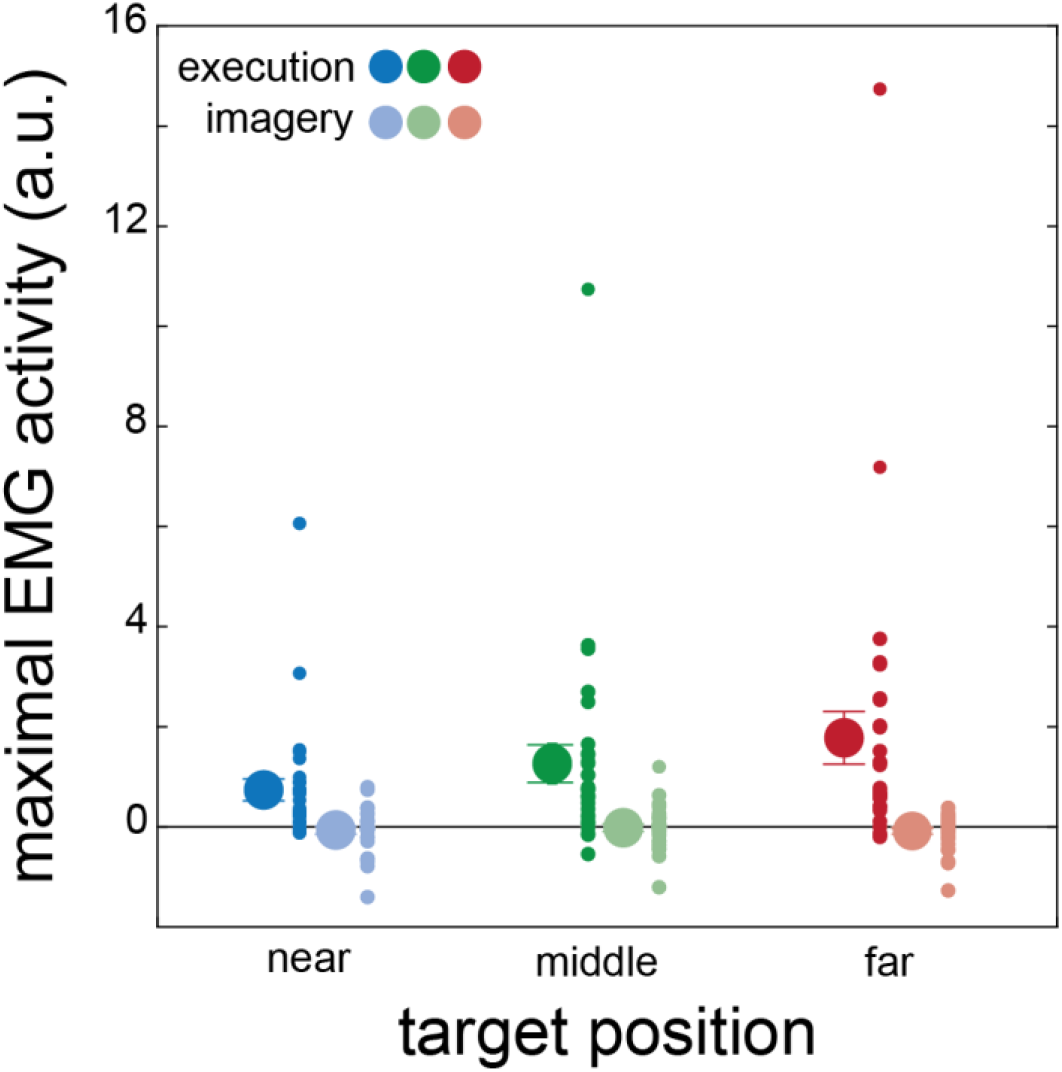
Muscle activity in Experiment 2. Peak muscle activity as a function of task and target distance relative to the resting EMG activity (= zero). Muscle activity was evident during motor execution but it was negligible during motor imagery. Averages with standard errors of the mean, together with individual participants’ data (small symbols) are shown.

Lastly, we explored whether the peak pupillary response was related to muscle activity. To this end, we ran two separate Pearson’s correlations between the maximal muscle activity and the peak pupil response, separately for the motor execution and motor imagery tasks. Neither of these tests revealed any systematic relationship (motor execution: r = −0.13, p = 0.219; motor imagery: r = −0.08, p = 0.406).

## Discussion

The main purpose of Experiment 2 was to rule out that pupillary responses related to motor imagery found in Experiment 1 were due to either the anticipated movement associated with the upcoming button press at the end of the trial, or spurious muscular activity during the imagery task. The results show that both motor execution and motor imagery led to significant pupillary dilations, independent of an anticipation of upcoming actions. Pupillary responses remained stronger during motor execution than motor imagery, just as in Experiment 1. This may be due to additional motor components associated with the overt reaching movement that was performed in the former but not in the latter condition. However, we did not find any correlation between the peak pupillary response and the peak muscle activity in the motor execution trials, which suggests that the strength of the pupillary dilation during these tasks is not necessarily affected by the underlying muscular activation.

## General Discussion

The goal of the present study was to investigate pupillary responses during motor execution and motor imagery of goal-directed reaching, as well as possible influences of target distance. Participants reached or imagined reaching to one of 3 targets that were placed at different distances from a start position. We replicate the typical mental chronometry findings (e.g., Papaxanthis et al., 2002) by demonstrating a strong and systematic correlation between actual and imagined movement times. Pupillary responses were greater during both motor execution and motor imagery relative to rest, but motor execution led to stronger responses than motor imagery. Task dynamics, varied by target distance, affected executed and imagined movement times; but they only had an influence on pupillary responses during motor execution and not during motor imagery. This might mean that pupillary responses during motor imagery do not necessarily reflect specific motor-related processes but that cognitive processes, such as recalling the target position or movement duration from memory, may also at least partially explain the dilations. In line with this notion, we also did not find systematic differences in pupillary responses between motor and non-motor imagery tasks, despite the fact that both types of imagery differed significantly from resting activity.

Pupillary responses were clearly pronounced during motor execution compared to the resting baseline, confirming previous work (e.g., Richer & Beatty, 1985; Rozado et al., 2017). Our findings further demonstrate that pupillary response characteristics during goal-directed reaching can be modulated by the distance to the target. For instance, the peak pupillary response was larger when participants reached to targets farther away, and this was systematic across both experiments. However, the time when this peak pupillary response occurred was not affected by target distance in either experiment. Thus, the reaching dynamics influenced only the maximal pupillary dilation but not its timing. This suggests that the effect of movement on pupil dilation was not caused by the longer movement times, in which case one might have expected peak pupillary responses to occur later in time. Since all movements employed here were single out-and-back reaches, it is possible that longer movement sequences could yield an effect on peak pupil dilation latency that was not visible in the current task.

Our study aimed to characterize the temporal dynamics of the pupillary response. Our time course analysis revealed that the onset and duration of pupil dilation was affected by the reaching distance to the target. More specifically, the further away one reaches, the earlier the pupil starts to dilate. This is evident not only when comparing the dilations to the resting baseline but also when comparing the responses among the different reaching targets. Because participants in both experiments were informed about the movement target before each trial, this difference in timing might reflect processes related to the planning and preparation of upcoming movements of different amplitudes (Moresi et al., 2008a, 2008b).

In line with previous work (O’Shea & Moran, 2016; Rozado et al., 2017), pupillary responses were also evident during motor imagery. However, we did not obtain any evidence that these responses reflect specific motor-related processes, as they were not modulated by movement parameters such as reaching amplitude. On the one hand, we found stronger peak pupillary responses during motor imagery than resting baseline. These dilations were evident over an extensive period of time, but lasted generally shorter than the responses during motor execution. On the other hand, pupillary responses were weaker during motor imagery than execution. Although these dilations could represent some motor-related processes, the current study did not find specific evidence to support this idea. For instance, the peak pupillary response during motor imagery was invariant across the 3 imagined reaching distances, suggesting that it is not modulated by movement complexity in the same way as the established pupil dilation during executed movements. The time course analysis also did not reveal any effects of target distance at any moment during the 5000 ms of recording. Please note that the target distance did affect the imagined *movement times*, similarly to how it affected the executed movement times.

The non-motor imagery task showed significant and prolonged pupil dilations, as reflected both in the peak pupillary response and in their temporal dynamics. These pupillary responses likely reflect general cognitive load or memory processes associated with recalling the previously shown painting, which is in line with other findings showing that cognitive tasks can cause the pupil to dilate (Hess & Polt, 1964; Kahneman & Beatty, 1966; Schuetz et al., 2021; Stoll et al., 2013). These pupillary responses were similar to those found during the motor imagery task. Given that the imagined reaching dynamics did not affect motor imagery pupillary responses, the combined results of motor and non-motor imagery tasks suggest that pupil dilation in the current imagery tasks likely reflects non-motor cognitive processes, at least for the case of simple, goal-directed reaching movements.

In Experiment 1, we asked participants to reach or to imagine reaching to different targets and then press a button once they had finished the task. This was important to assess the modulation of movement times by the task and the target distance, and to relate our findings to previous work that used the mental chronometry approach (e.g., Papaxanthis et al., 2002; Munzert et al., 2015). However, we wanted to rule out that this anticipated manual task of pressing the button might have caused an anticipatory pupillary dilation that could at least partially explain the results of Experiment 1 (Richer & Beatty, 1985). We therefore addressed this in Experiment 2, by asking a different set of participants to reach or imagine reaching to the same targets as before but without pressing a button at the end of the trial. The respective results were highly similar to those obtained in Experiment 1, suggesting that pupillary responses during motor execution and motor imagery in Experiment 1 are unlikely to be caused by preparatory activity related to an anticipated manual action.

The pupillary responses during motor execution most likely reflect motor processes associated with the ongoing movement. Previous work has already shown that pupil dilation is stronger when performing more complex movements (Richer & Beatty, 1985) or when an action requires greater precision (Fletcher et al., 2017). However, it is not yet clear what aspect of these factors (i.e. movement complexity or required precision) might influence the pupillary response. We therefore assessed the EMG activity during reaching to the same targets in Experiment 2. Although we found stronger maximal EMG activity when reaching to farther away targets, we did not find any relationship between the strength of the maximal EMG activity and the peak pupillary response. Therefore, the modulation of the pupil dilations by motor execution and by target distance that we report is unlikely caused by the underlying muscle activity. Similarly, EMG activity was negligible during motor imagery as compared to the resting baseline, suggesting that pupillary responses also during motor imagery are unlikely to be caused by underlying EMG activity.

In sum, our results show that pupillary responses during motor execution of goal-directed reaching movements are modulated by the dynamics of the movement, such as movement amplitude, and that this modulation seems independent of the associated muscular activity. We further show that motor imagery of the same goal-directed movements leads to systematic pupillary dilations, which are however independent of the dynamics of the imagined movement. Rather, they are more similar to those pupillary responses elicited during a non-motor imagery task, suggesting that pupillary responses during motor imagery reflect general cognitive processes related to motor imagery and not necessarily the underlying motor processes.

## Acknowledgments

We would like to thank Nora Jaeger for her reliable support with data collection. KF and DV are supported by the German Research Foundation (DFG) - Collaborative Research Center SFB/TRR135, project A4, under grant agreement 222641018. KF is also supported by “The Adaptive Mind” funded by the Excellence Program of the Hessian Ministry for Higher Education, Research, Science and the Arts. DV is also supported by the German Research Foundation (DFG) under grant agreement VO 2542/1-1. Raw data from the reported experiments can be found at https://osf.io/ugcn8/.

